# Marker-assisted introgression of multiple resistance genes confers broad spectrum resistance against bacterial leaf blight and blast diseases in Putra-1 rice variety

**DOI:** 10.1101/750216

**Authors:** Samuel C. Chukwu, Mohd Y. Rafii, Shairul I. Ramlee, Siti I. Ismail, Yusuff Oladosu, Isma’ila Muhammad

## Abstract

This experiment was conducted with the aim of introgressing multiple resistance genes against bacterial leaf blight (BLB) and blast diseases through marker-assisted backcross breeding. Two dominant (*Xa4* and *Xa21*) and two recessive (*xa5* and *xa13*) BLB resistance genes were introgressed into a Malaysian elite rice variety Putra-1 with genetic background of three blast resistance (*Piz, Pi2* and *Pi9*) genes and high yielding. Eight polymorphic tightly linked functional and SSR markers were used for foreground selection of target genes. 79 polymorphic SSR markers were used in background selection. The plants were challenged at initial stage of breeding and challenged again at BC_2_F_2_ with the most virulent Malaysian pathotypes of *Xoo* (P7.7) and *Magnaporthe oryzae* (P7.2) to test their resistance. Results obtained from foreground marker analysis showed that the BC_1_F_1_ and BC_2_F_1_ both fitted into the Mendel’s single gene segregation ratio of 1:1 for both *Xoo* and blast resistance. At BC_2_F_2_, result obtained indicated that foreground marker segregation fitted into the expected Mendelian ratio of 1:2:1 for blast resistance only. Marker-assisted background selection revealed high percentage of recurrent parent genome recovery (95.9%). It was concluded that resistance to *Xoo* pathotype P7.7 in IRBB60 was neither due to two independent gene action nor epistasis but substantially due to single nuclear gene action. Also, the inheritance of blast resistance in the pyramided lines to pathotype P7.2 was also attributed to single gene action. The incorporation of four bacterial leaf blight and three blast resistance genes (*Xa4+xa5*+*xa13*+*Xa21+Pi9+Pi2*+*Piz*) in the newly developed lines provides for broad spectrum and durable resistance against the two major diseases studied.

## Introduction

Rice production is a very important activity all around the world. Many African and Asian countries rely on rice as a source of income and daily calories [1, 2, 3]. Bacteria leaf blight (BLB) which is caused by the pathogen *Xanthomonas oryzae* pv*. oryzae* is one of the most serious diseases responsible for significant yield reduction in rice. The disease constraints the photosynthetic area of the crop thereby causing partial grain filling and this leads to poor yield [4]. Use of resistant varieties of rice is the best approach to guard against the disease as chemical control is not effective. To date, 42 *Xoo* resistance genes have been identified in rice and some of them have been incorporated into new varieties [5]. Generally, the disease is favoured by temperatures at 25 to 34°C, with relative humidity more than 70% and most rice growing regions in Malaysia are affected by the disease [6]. Host plant resistance offers the most viable, low cost effective and sustainable strategy to manage the disease than chemical control and biological method. Use of multiple R-genes is the most attractive approach to combat BLB [7].

Rice blast caused by the fungus *Pyricularia oryzae* [*Magnaporthe oryzae* (anamorph)] is one of the biotic agents responsible for severe yield loss in rice [8]. It is among the most destructive diseases affecting rice in Malaysia and other parts of the world’s major rice growing regions. The disease causes considerably large grain yield losses in almost all major ecosystems where it is grown [9, 10]. Blast infection could be noticed in every stage of rice growth and the symptoms occur in all above-ground parts. The early symptoms look like white/gray/green spots or necrosis surrounded by dark green colour. Molecular markers that co-segregate with the gene are a powerful method to develop a resistant cultivar. The availability of different molecular markers allows characterization of genes of interest [11].

Resistance genes for BLB and blast are usually obtained from rice varieties, land races, primitive cultivars and wild species. About 100 blast resistance genes have been identified, and many of them have been cloned and characterized [12]. Variability in pathogens leading to overcoming of resistance effect of major gene after 4–5 years, has forced the breeder for continuous search of novel R-genes. Widespread use of resistant cultivars leads to development of selection pressure in the pathogen that resulted into breaking of resistance barrier. Majority of R-genes are dominant in nature, but recessive gene has also been reported [6]. Dominant R-genes for BLB are *xa-1, xa-3, xa-4, xa-7, xa-10, xa-14, xa-21* and *xa-22(t)* and some major recessive genes are *xa-5, xa-13*. Most blast R-genes are dominant in nature. Pyramiding of genes is an effective strategy for enhancing the durability of R-genes over time and space. Marker assisted selection is one of the most sustainable and attractive technology that facilitates incorporation of two or more major genes in different genetic background of rice. In the presence of a number of virulent pathogens, the genes may differ in their level of resistance to them. Thus, gene pyramiding is currently being pursued in an effort to develop more durable and comprehensive resistance rice varieties to combat the effect of BLB and blast pathogens. Therefore, this work was carried out to introgress resistance genes from two genetically diverse parents using marker-assisted selection for broad spectrum resistance against BLB and blast diseases.

## Materials and Methods

### Genetic materials and method of plant breeding

The donor parent IRBB60 which contains four BLB R-genes *Xa4, xa5, xa13* and *Xa21* [6] was collected from the International Rice Research Institute (IRRI), the Philippines. *Xa4* and *Xa21* are the dominant genes while *xa5* and *xa13* are in the recessive condition. The recurrent parent was Putra-1, a high yielding rice variety that is resistant to blast disease with genetic background of three blast R-genes, namely; *Piz, Pi2, Pi9*. Putra-1 variety was developed at the Institute of Tropical Agriculture and Food Security (ITAFoS), Universiti Putra Malaysia, from a cross between MR219 × Pongsu Seribu 1 [13]. Putra-1 was used as the female parent to hybridize the male parent IRBB60 and F_1_ plants were developed. The Putra-1 was subsequently used as the recurrent parent during backcrossing stages to backcross with F_1_ leading to development of BC_1_ and later backcrossed to BC_1_ leading to development of BC_2_ plants. Marker-assisted backcross breeding method was adopted up to BC_2_F_2_ generation. At each stage of backcrossing, at least 200 lines were genotyped using polymorphic tightly linked foreground markers for selection of target genes. At BC_1_F_1_ and BC_2_F_1_, only plants that showed heterozygous alleles for both bacterial leaf blight and blast resistance genes were selected while plants that showed homozygous alleles for both resistances genes at BC_2_F_2_ were finally selected. Foreground selection for multiple target genes was done up till BC_2_F_2_ where recombined and pure homozygous lines for all seven target genes were selected. Background selection using 79 polymorphic SSR markers was also done up till BC_2_F_2_ to ensure that only lines with recovered recurrent parent genome as well as pyramided seven target genes for BLB and blast were selected. Phenotypic selection was combined with foreground and background selections at every generation to reconfirm the results of molecular screening. Management practices such as planting spacing, weed control, fertilizer application, pest and disease control were done according to the Malaysian Agricultural Research and Development Institute (MARDI) recommended manual.

### Molecular marker screening

Fifteen molecular markers including both SSR and functional markers found in literature that were linked to *Xoo* resistance genes were screened to confirm their polymorphism between the two parents; Putra-1 and IRBB60. Also, two SSR DNA-based markers found in literature that were linked to blast resistance genes were also tested for their polymorphism between the two parents. A total of 472 rice SSR markers were screened to select the polymorphic markers between the parents Putra-1 and IRBB60 which were subsequently used for background selection.

### Genomic DNA extraction

To extract genomic DNA, young and fresh rice leaves were collected from 4-week-old individual plants using the CTAB (Cetyltrimethylammonium bromide) method modified from Ashkani et al. (2013).

### Confirmation of DNA bands

Three μl of the extracted DNA were mixed with 2 μl of DNA loading dye on top of a parafilm wax and then loaded into the wells of a 3g metaphor agarose gel and subjected to electrophoresis. After running the electrophoresis for about 45 minutes at 90V, 2500 aM, the gel was viewed on image viewer machine for confirmation that the DNA bands were present.

### DNA identification and quantification

One μl of each DNA sample was put on NanoDrop spectrophotometry (ND-1000, NanoDrop Technologies Inc., Wilmington, DE, USA) and relative purity with concentration of the extracted DNA was estimated from the computer display data value. The final concentration of each DNA sample was diluted with TE buffer (10 mMTris-HCl, pH 8.0, 1 mM EDTA, pH 8.0) to get the required concentration and maintained in a refrigerator of −20 °C for PCR analysis.

### Polymerase chain reaction (PCR)

After confirming the DNA quantity and quality, the selected suitable samples were taken for PCR analysis using 15 μl PCR mixture. Thereafter, the PCR tubes were loaded in the PCR machine (Product specification: Thermocycler, T100TM, Bio-Rad, UK) and the PCR was run following three hours plus three minutes touchdown PCR programme. After initial denaturation for 5 min at 94 °C, each cycle comprised 1 min denaturation at 94 °C, 1 min annealing at 55 °C, and 2 min extension at 72 °C with a final extension for 5 min at 72 °C at the end of 35 cycles.

### Gel Electrophoresis

A 3% Metaphor^TM^ agarose (Lonza) gel was prepared using 3 g Metaphor^TM^ agarose powder dissolved in 100 ml 1×TBE buffer with the aid of the microwave machine. After proper dissolution, 10 ul of Gel view was added and then shaked to mix very well. The gel was run at a steady voltage of 90 set at maximum 2500A for 45 min and the resultant bands were visualized using Molecular Imager® (Product specification: GelDocTM XR, Bio-Rad Laboratories Inc., USA).

### DNA/Marker scoring

Marker scoring was done by scoring the DNA bands obtained from the computer display result of Gel document (Image viewer). Single bands were scored either as homozygous allele similar to the recipient (recurrent) parent/female represented with the letter ‘A’ or homozygous allele similar to the donor (male) parent represented with the letter ‘B’. Double bands were scored as heterozygous allele (heterozygote) carrying the genes of both parents and represented with the letter ‘H’.

### Disease challenging for bacterial leaf blight resistance

Diseased samples of rice were collected from 15 different locations used for isolation of *Xoo*. Infected leaf pieces 28 × 7 mm of rice were excised with sterile scalpel. The leaf surface was sterilized with 1% Clorox (Sodium hypochloride) for three minutes and then washed with sterile distilled water (SDW) twice of one to two minutes each. The cut leaf pieces (6-7) were transferred to nutrient agar (NA) medium after drying on sterile blotting paper and incubated at room temperature (25-28 °C) for 72 hrs [14]. The single bacterial colonies were sub-cultured onto NA plates and to get pure culture, cultures were preserved for longer duration (at 4 °C) on NA slants. The isolates were subjected to the pathogeniecity test to confirm the pathogenicity of the isolated bacteria [15]. Pure cultures of bacterial colonies were suspended into 5 ml distilled water and the concentration of inoculums was adjusted to 10^8^ cfu/ml [9] and homogenised. The plants were sprayed with water to create wet conditions favourable for disease development. Inoculation was done by cutting five leaves, approximately 5 cm from the tips of each line with sterilized scissors dipped in the prepared inoculum. After 14 days, disease data were assessed to identify the degree of pathogenicity on 0 to 9 disease rating scale using Standard Evaluation System of IRRI [16]. The disease incidence and severity were determined. At BC_2_F_2_, pure cultures of the most virulent *Xoo* pathotype P7.7 obtained from MARDI was used for inoculation.

### Disease challenging for blast resistance

On a special request to the Malaysia Agriculture and Research Development Institute (MARDI), Seberang Perai, the most virulent pathotype P7.2 of blast pathogen *Magnaporthe oryza* was provided. Potato dextrose agar (PDA) was prepared in the Physiology laboratory, ITAFoS and used for sub-culturing the collected pathotype P7.2. Plating was done under a sterile condition in the laminar flow hood and the petri dish was allowed to solidify and sealed with parafilm to avoid any form of contamination. The plate cultures were streaked from the master culture using a sterilized needle and sealed with tape to avoid contamination [17]. The plates streaked with pathogen were kept under a light and dark cycle of 16 and eight hours, respectively, under room temperature for four days. Then the cultures were incubated for another 10 days at 28 °C for maximum conidial production. A sterile slide was used to harvest the conidia by scraping the fungal mycelia on the culture plates. Then, the conidia suspension was filtered using the cheese cloth to remove the mycelia and agar pieces [15, 18]. In order to measure the concentration of the initial conidial suspension, a conidial suspension was delivered to both chambers of haemocytometer by capillary action [19, 20]. The spore concentration was adjusted to 1.5 x 10^5^ spores/ml standard by haemocytometer. Prior to inoculation, 0.05% Tween 20 was added to the suspension to increase the spores adhesion on leaves of plants. The donor parent (IRBB60), recurrent parent (Putra-1) and the newly developed lines were inoculated with the pathogen. In order to maintain an ideal environment for the disease development, the plants were covered with plastic bags, maintaining about 90% humidity. After nine days of inoculation, the challenged plants were phenotypically screened for blast disease symptoms. The disease incidence and severity were determined [16].

### Characterization for agro-morphological traits

The agro-morphological performance was evaluated using lines that already recovered their recurrent parent genome with high yielding genetic background and all seven target genes pyramided. Agronomic traits associated with yield and yield components were measured and recorded. These included number of days to 50% flowering and maturity, plant height, number panicles per plants, effective tillers per plants, panicle length, number of filled grain per plant, 1000-grain weight, total yield per plant, seed length, seed width, flag leaf length, flag leaf width, leaf area, leaf area index, and other traits of economic importance. These traits were recorded from all of the best selected lines of BC_2_F_2_ along with the recurrent parent. The morphological and agronomical characteristics were measured following standard procedures.

### Statistical analysis

The genotypic data obtained from the DNA scoring were subjected to Chi-square analysis using the formular:

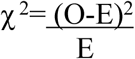

Where: O and E are the observed and expected values. The χ^2^ calculated was compared with the χ^2^ tabulated at P=0.05 in order to determine whether the markers showed significant differences from the Mendelian segregation ratio of 1:1 between the observed and expected values obtained from the backcross population. The DNA scoring data obtained from background selection with the aid of the 79 polymorphic markers were further subjected to analysis using the genetic software Graphical Genotyper (GGT 2.0) [21]. This analysis was done to determine the genotypes with maximum recurrent parent genome recovery. With respect to selection for maximum recovery, the sorted SSR data were analysed with the Graphical Genotyper (GGT 2.0) software.

## Results and Discussion

### Marker assisted Pyramiding of bacterial leaf blight and blast resistance genes

The result obtained on initial parental polymorphism survey showed a clear polymorphism between the two parents Putra-1 (recurrent) × IRBB60 (donor) (Figure 1). The parental polymorphism was identified between the two parents in all seven target resistance genes for both BLB and blast using Xa21FR and pTA248 (*Xa21*), Xa13prom (*xa13*), RM21 and RM13 (*xa5*), MP (*Xa4*), RM8225 (*Piz*) and RM6836 (*Pi9*, *Pi2* and *Piz*) (Table 1).

**Figure 1:**
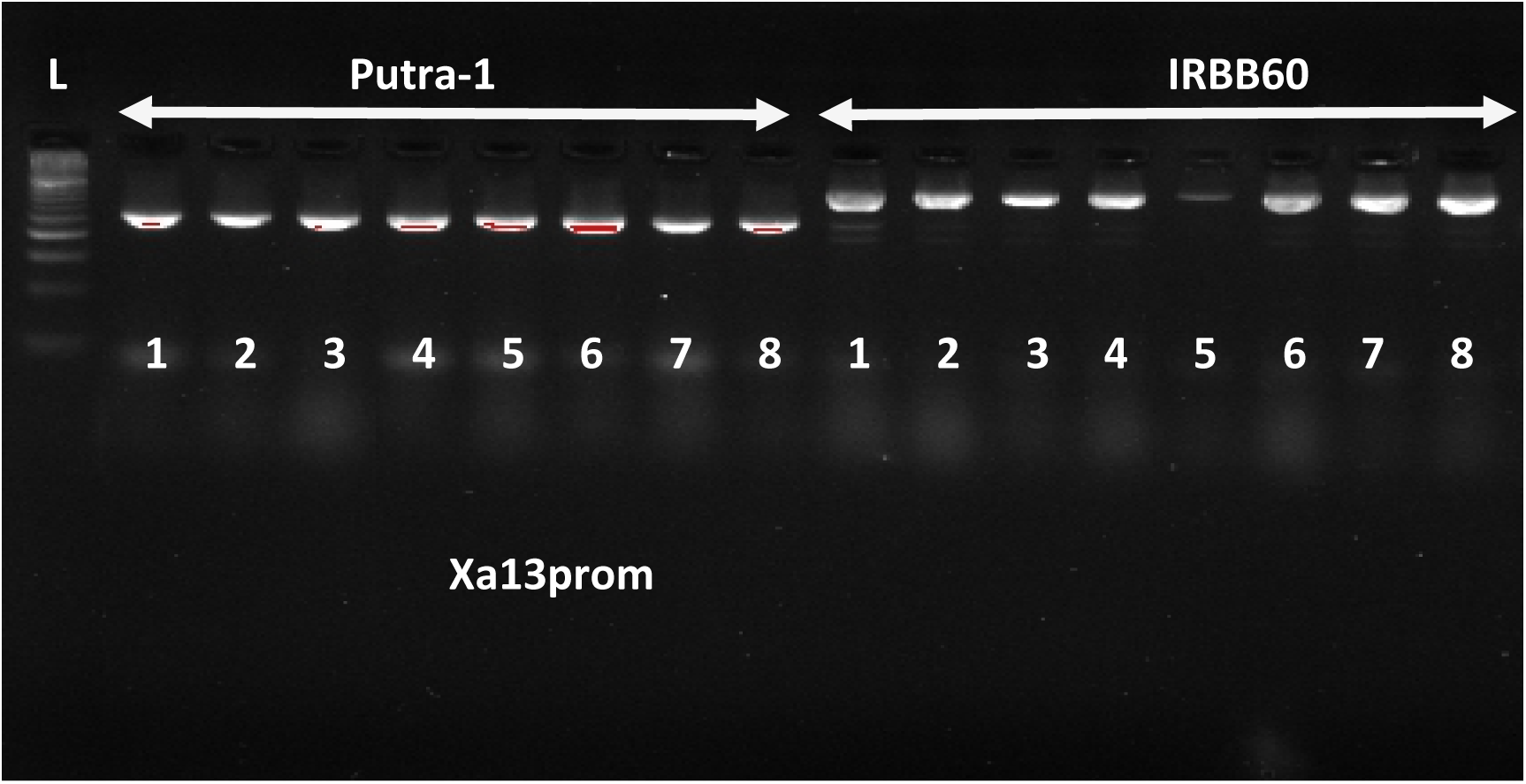
Molecular screening of parental genotypes for polymorphism using Xa13prom foreground marker linked to the *xa13* gene.

**Table 1:**
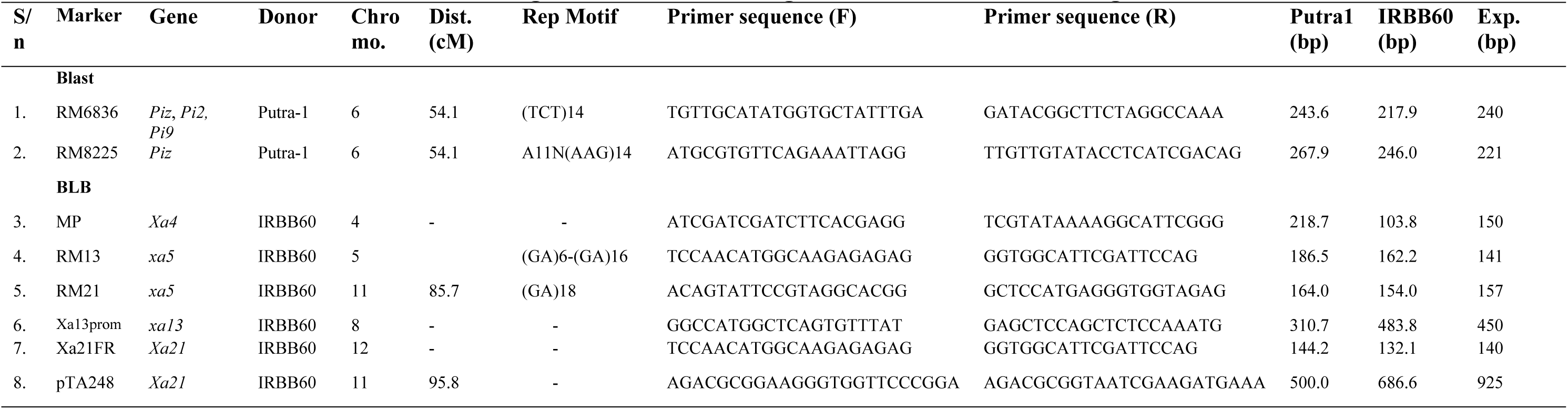
Information of markers used for foreground selection of target BLB and blast resistance genes

### F_1_ confirmation using polymorphic tightly linked foreground markers

A cross was made between Putra-1 and IRBB60 which produced the F_1_ seeds. A total of 72 F_1_ plants were grown from the F_1_ seeds and tested for heterozygosity using the foreground polymorphic markers. Four BLB (Xa21FR, Xa13prom, pTA248 and RM13) and two blast (RM8225 and RM6865) polymorphic foreground markers were used for the foreground selection. Out of the 72 grown F_1_ plants, a total of 46 plants were confirmed heterozygous true F_1_ plants and scored as ‘H’ using the Xa21FR polymorphic tightly-linked foreground marker. In order to determine the plants that carry all the target genes, the 46 heterozygous plants were screened with the other foreground markers and five plants were confirmed to carry both BLB (*Xa4, xa5, xa13* and *Xa21*) and blast (*Piz*, *Pi2* and *Pi9*) R-genes. These five progenies were selected for use in crossing to produce BC_1_F_1_ because the *Xoo* and blast resistance genes have been confirmed to be pyramided into them. Figure 2 shows banding patterns obtained from genotyping of F_1_ plants.

**Figure 2:**
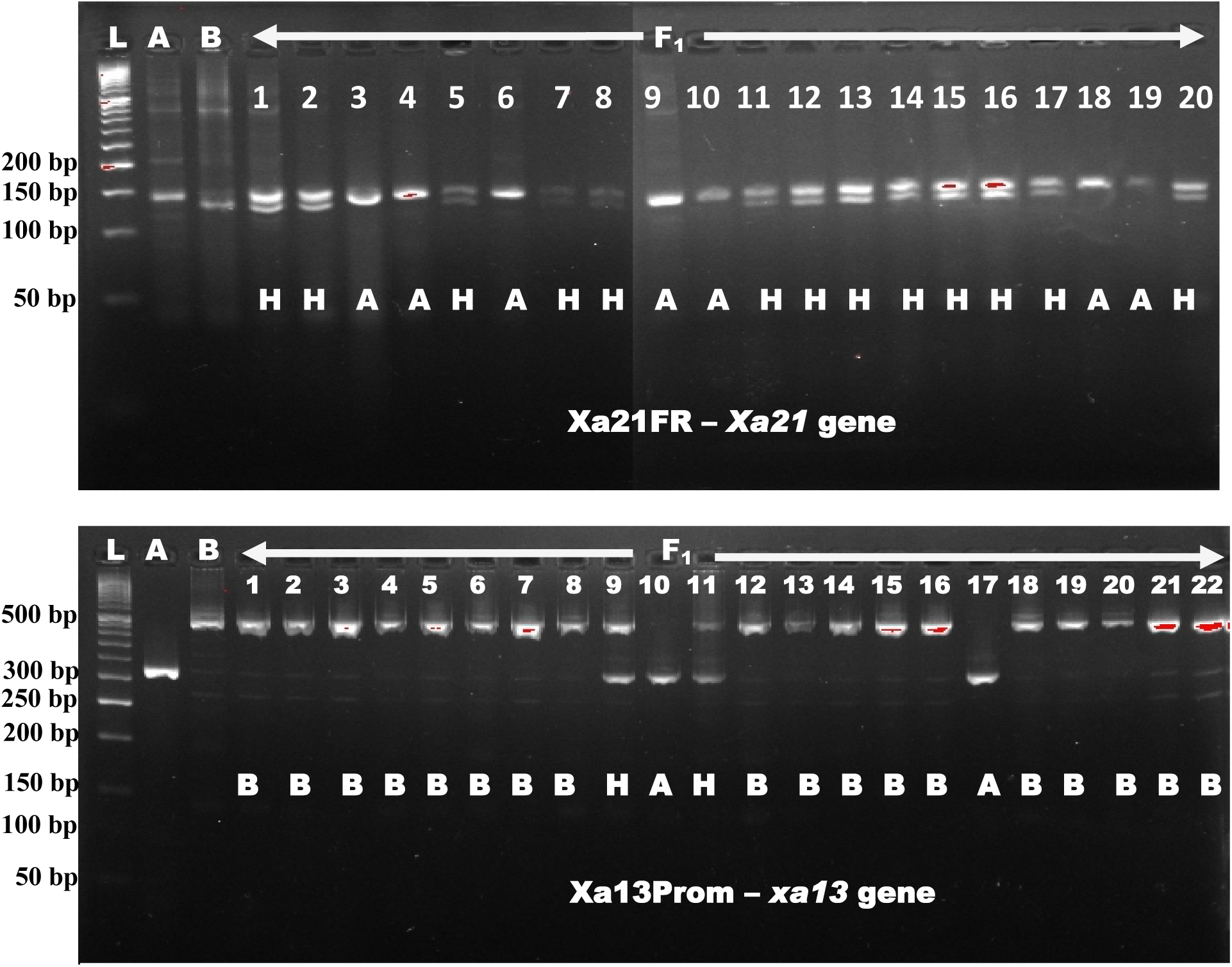
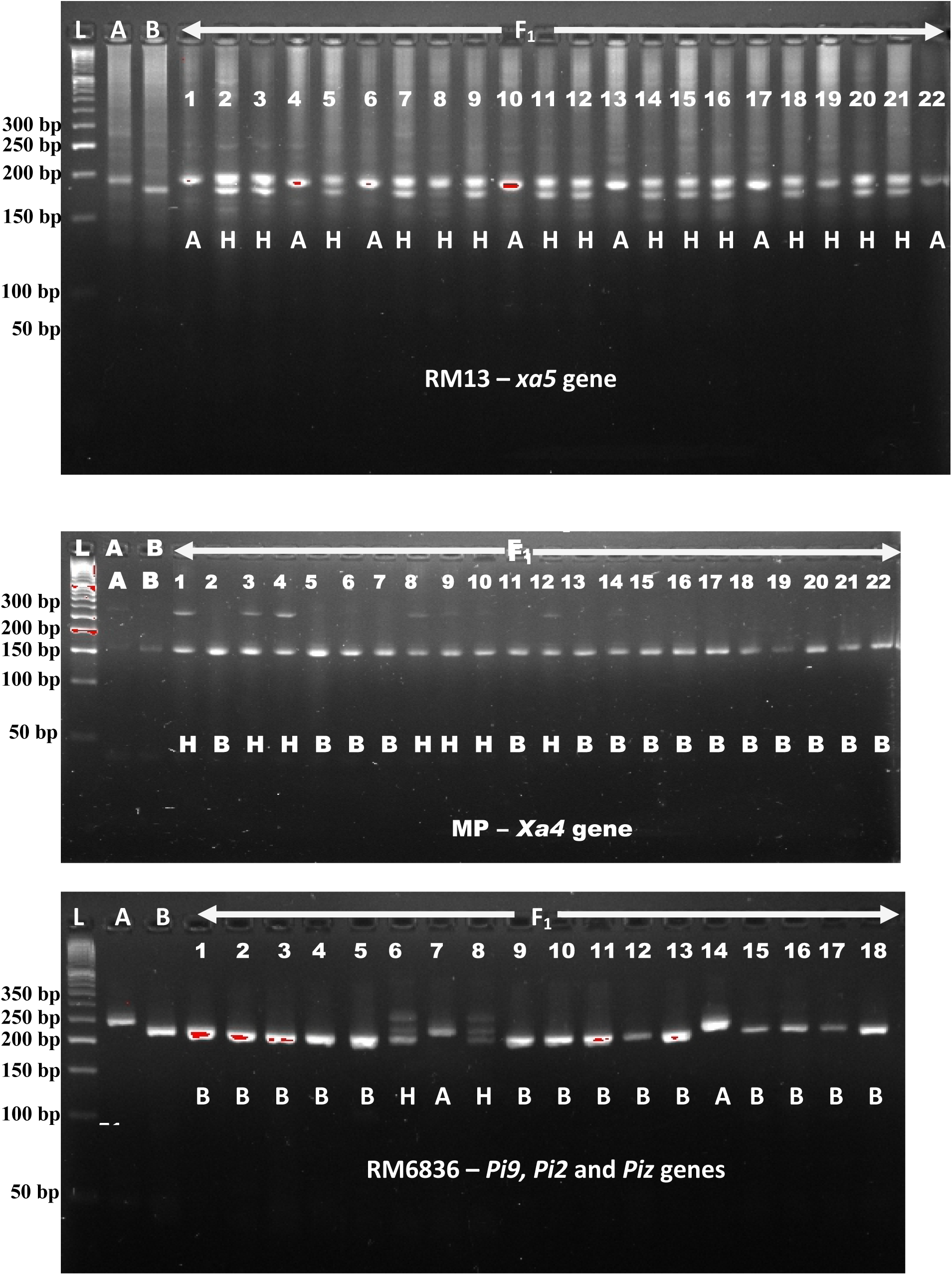
Genotyping and foreground marker-assisted selection of F_1_ progenies using polymorphic tightly linked foreground markers for pyramiding of BLB and blast target R-genes. A: Putra-1; B: IRBB60. H: heterozygote. 1, 2, 3, … 22 are some F_1_ plants. Electrophoresis running on 3% metaphor agarose gel stained with 10ul of gel view per 100 ml. L: 50bp DNA ladder.

### Marker genotyping of BC_1_F_1_ population

A total of 288 BC_1_F_1_ progenies were obtained from a cross between the best F_1_ progeny (F_1_ with both BLB and blast R-genes) and Putra-1 variety as female and male parents respectively. Out of the 288 BC_1_F_1_ progenies, 108 plants were heterozygous for *Xa21* gene (144.2 bp and 132.1 bp), 125 for *xa13* gene (310.7 bp and 483.8 bp), 118 for *xa5* (186.5 bp and 162.2 bp), 122 for *Xa4* (218.7 bp and 103.8 bp) and 112 for blast R-genes *Pi9*, *Pi2* and *Piz* (243.6 bp and 217.9 bp). The Chi-square result of the six foreground markers showed no significant difference in the BC_1_F_1_ progenies from a 1:1 Mendelian segregation ratio. The result indicates that the foreground markers segregated according to the 1:1 segregation ratio for single gene model. Out of all heterozygous BC_1_F_1_ plants, the result obtained from foreground marker-assisted genotyping showed that only nine plants carried all the target genes of BLB and blast resistance. The nine selected plants were later subjected to background selection. Figure 3 shows some banding patterns of BC_1_F_1_ foreground selection.

**Figure 3:**
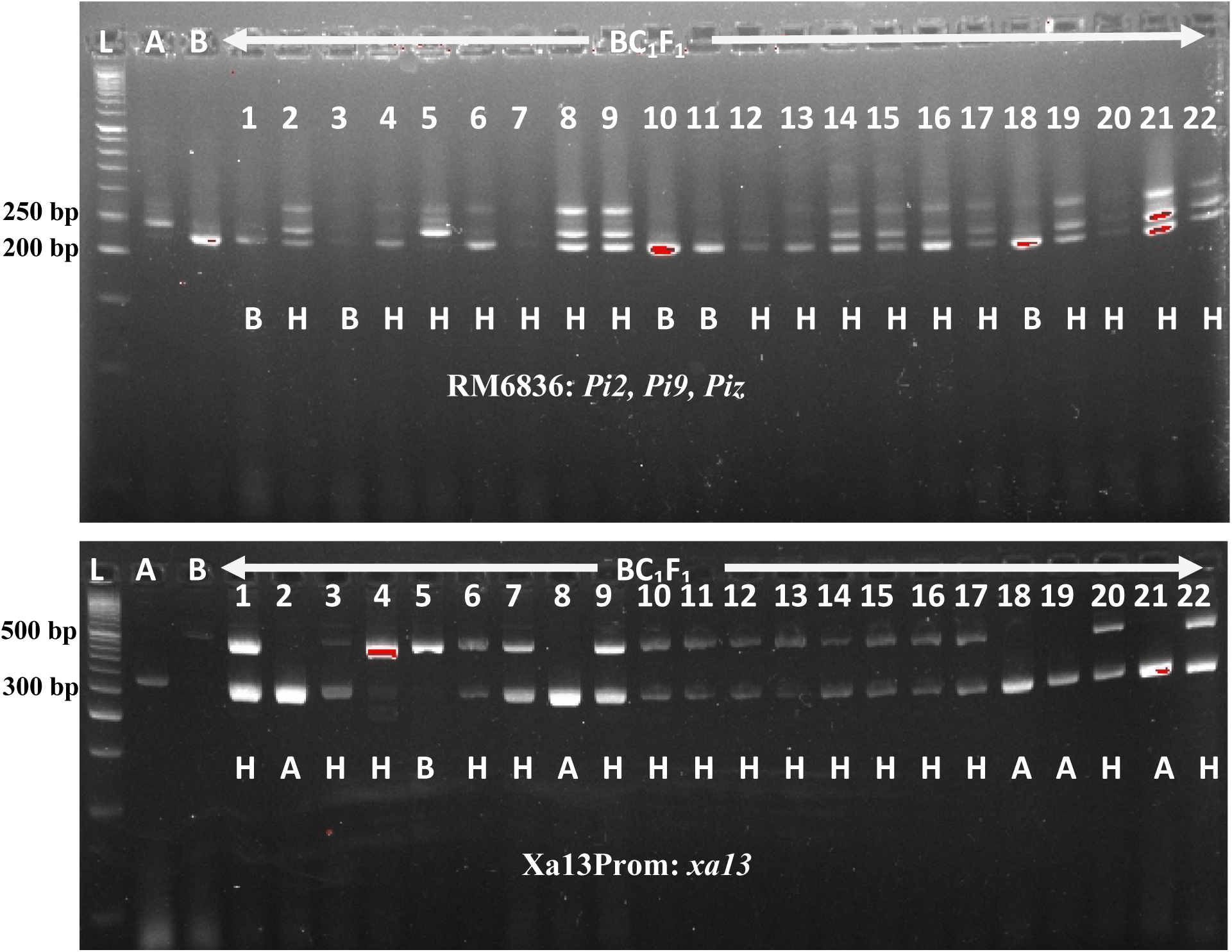
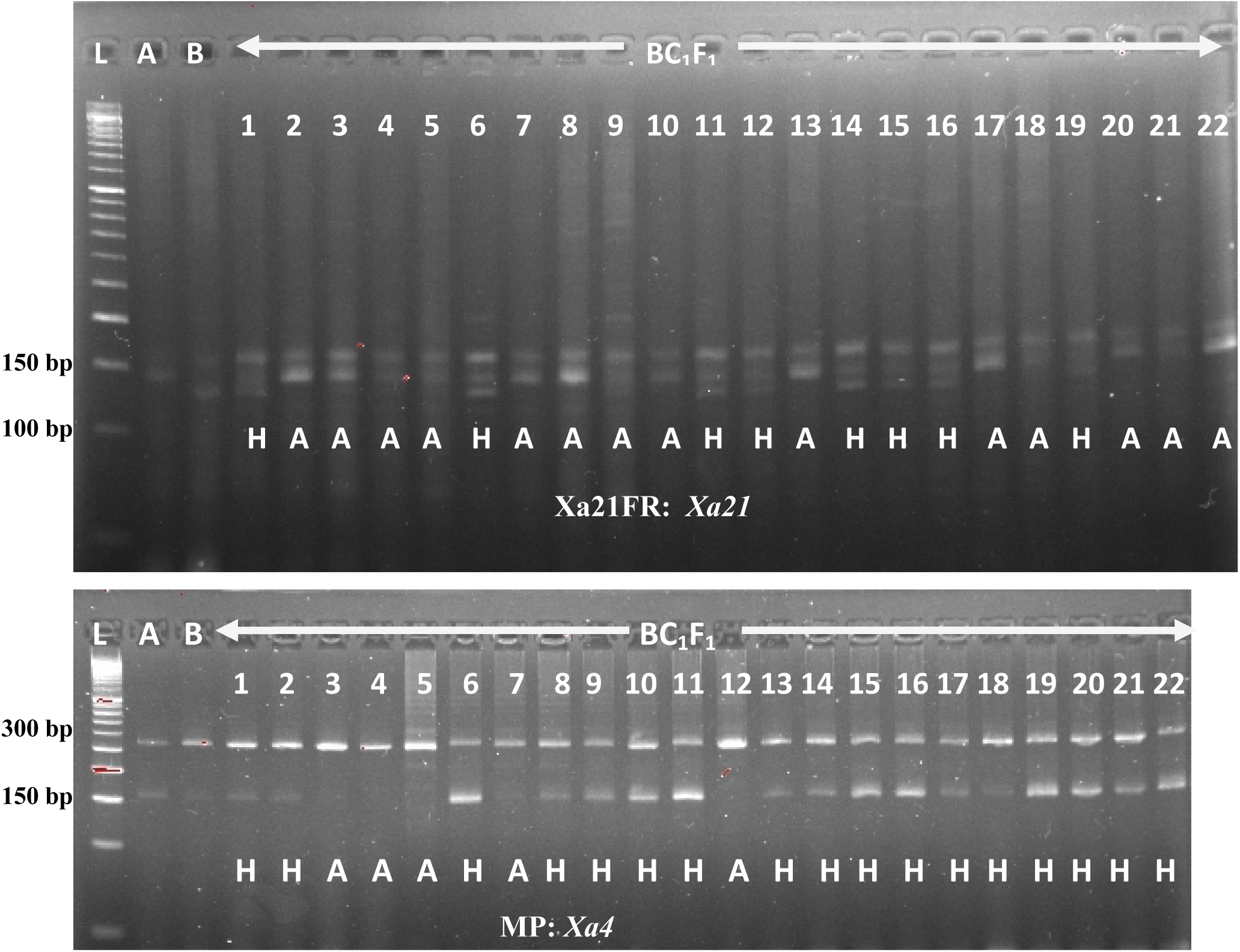
Genotyping and foreground marker-assisted selection of BC_1_F_1_ progenies using polymorphic tightly linked foreground markers for pyramiding of BLB and blast target R-genes. A: Putra-1; B: IRBB60. H: heterozygote. 1, 2, 3, … 22 are some BC2F1 plants. Electrophoresis running on 3% metaphor agarose gel stained with 10ul of gel view per 100 ml. L: 50bp DNA ladder

### Marker genotyping of BC_2_F_1_ population

At BC_2_F_1_ generation, a total of 268 rice crops were grown from the seeds of the six best BC_1_F_1_ plants already confirmed and selected. The pyramiding of four target BLB and three specific blast resistance genes were confirmed in 14 BC_2_F_1_ progenies with the aid of the foreground markers. The gel images of some of the BC_2_F_1_ progenies as extracted from the computer displayed image viewer document are as presented in Figures 4. Out of the 268 BC_2_F_1_ progenies, 106 plants each were heterozygous for *Xa21* (144.2 bp and 132.1 bp) and *xa13* (310.7 bp and 483.8 bp) genes, 104 for *xa5* (164.0 bp and 154.0 bp), 96 for *Xa4* (218.7 bp and 103.8 bp) and 104 for blast R-genes *Pi9*, *Pi2* and *Piz* (243.6 bp and 217.9 bp). The Chi-square result obtained indicated a non-significant difference and/or goodness of fit to the 1:1 Mendelian expected test cross ratio for a single gene model at BC_2_F_1_. This is an indication of close association with the BLB and blast genes for resistance to these particular bacterial and fungal infections. The result confirms the successful pyramiding of the target genes into the BC_2_F_1_ populations using marker-assisted foreground selection. Foreground markers are closely associated or linked to the gene of interest on the same chromosome and as such are carried from one generation to another [22, 23].

**Figure 4:**
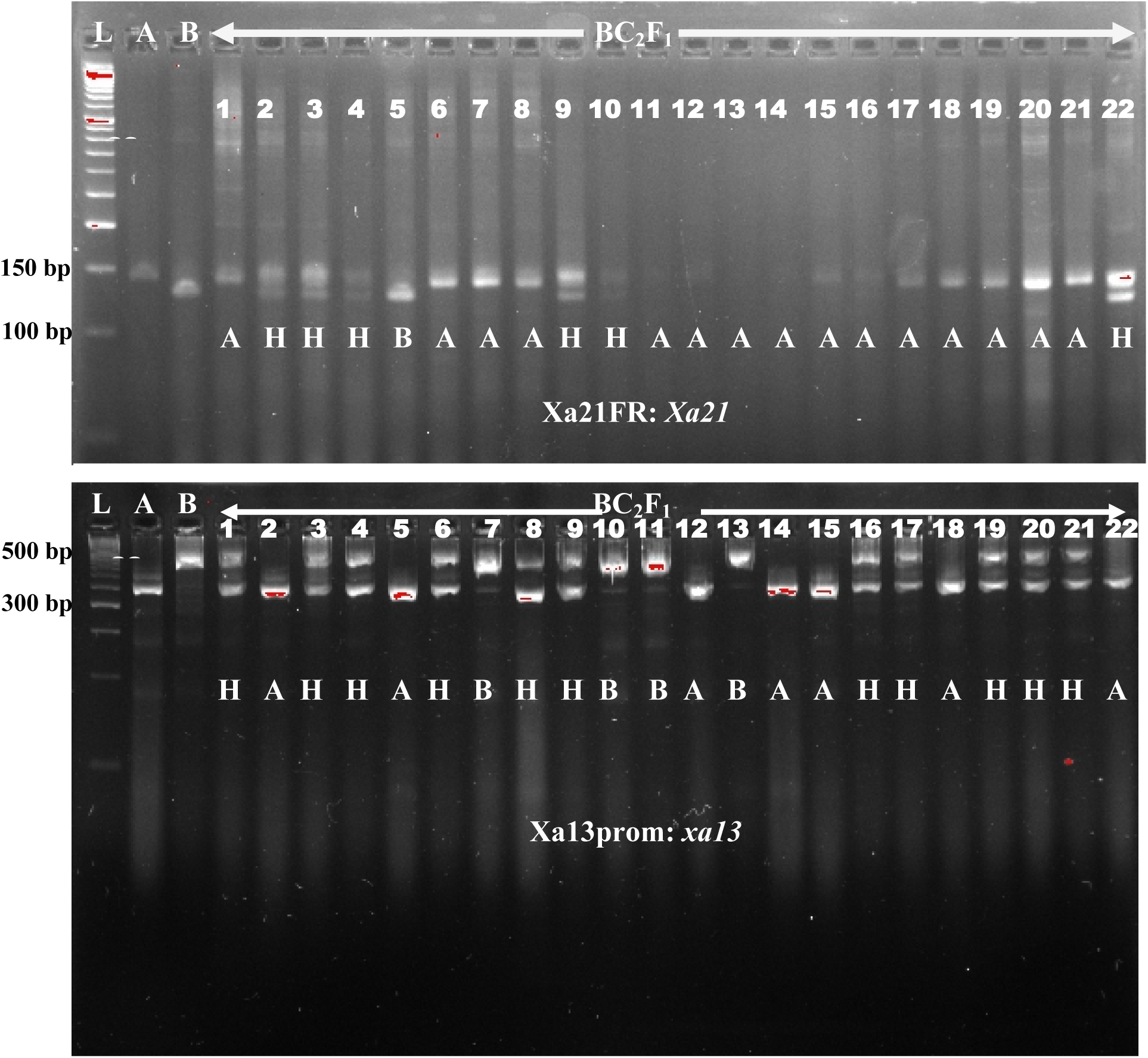
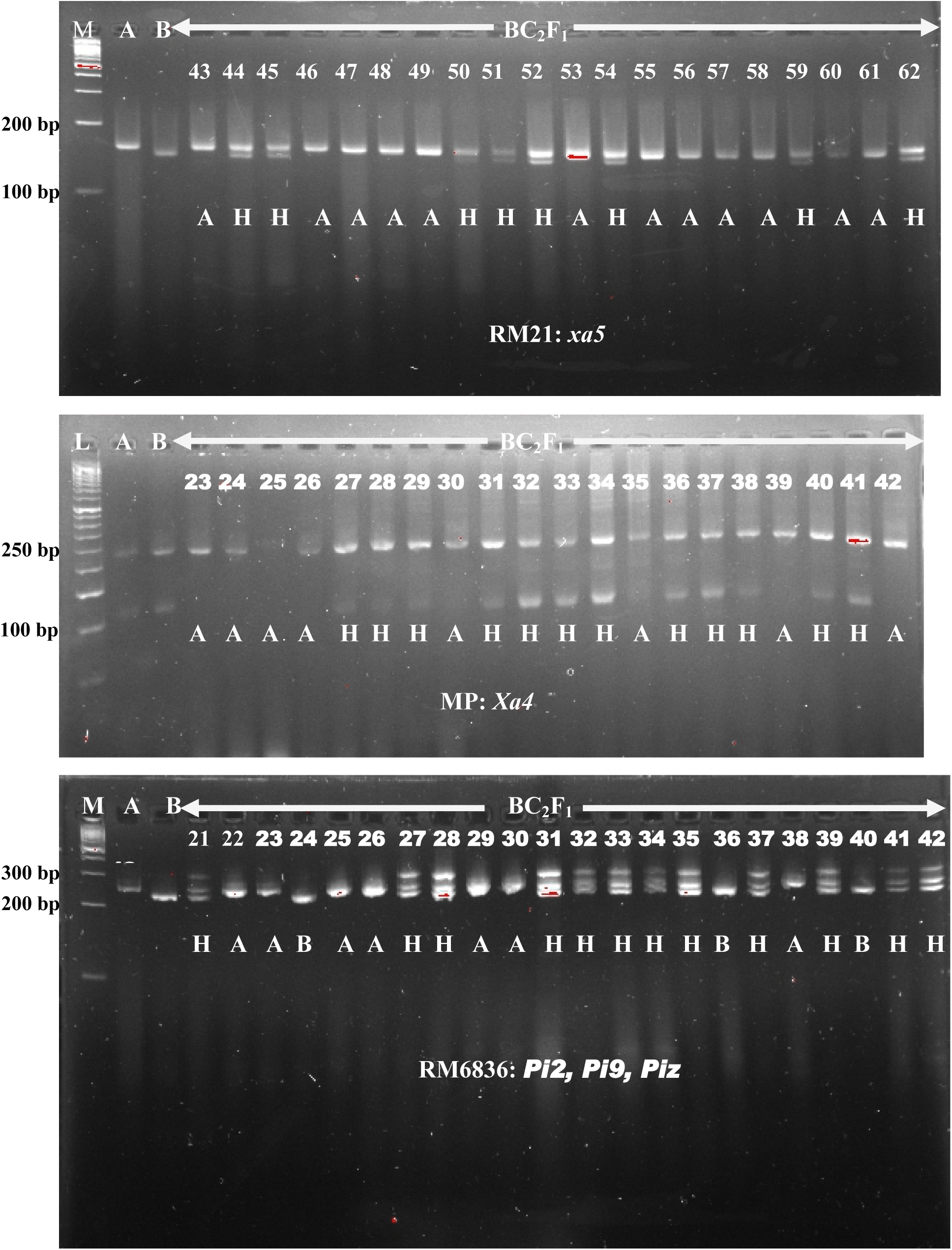
Genotyping and foreground marker-assisted selection of BC_2_F_1_ progenies using polymorphic tightly linked foreground markers for pyramiding of BLB and blast target R-genes. A: Putra-1; B: IRBB60. H: heterozygote. 1, 2, 3, … 42 are some BC_2_F_1_ plants. Electrophoresis running on 3% metaphor agarose gel stained with 10ul of gel view per 100 ml. L: 50bp DNA ladder, M: 100bp DNA ladder

### Marker genotyping of BC_2_F_2_ population

The result obtained on foreground selection of BC_2_F_2_ population indicated that out of 218 BC_2_F_2_ progenies screened, 17 plants had homozygous allele for *Xa21* gene (132.1 bp), 32 had homozygous allele for *xa13* gene (483.8 bp), 29 had homozygous allele for *xa5* gene (154.0 bp) while 115 plants had homozygous allele for *Xa4 Xoo* R-gene (103.8 bp). The result from Chi-square analysis indicates that the marker segregation ratios obtained for *Xoo* resistance did not correspond with the expected 1:2:1 Mendelian segregation ratio. However, at earlier backcrossing stages of BC_1_F_1_ and BC_2_F_1,_ the foreground marker segregation ratios agreed with the expected Mendelian segregation ratio of 1:1 for single genes. The marker segregation skewed towards the homozygous alleles (A) of Puta-1 susceptible to the bacterial leaf blight infection but resistant to blast fungal disease. Also, the result showed that out of the 218 BC_2_F_2_ lines selected, 69 had homozygous blast R-gene alleles (243.6 bp) similar to Putra-1, 100 progenies displayed heterozygous bands for both blast and *Xoo* R-genes. The other 49 progenies had homozygous *Xoo* R-gene alleles similar to IRBB60. Unlike the result obtained in *Xoo* resistance marker segregation analysis, the Chi-square result for blast resistance marker segregation showed that the marker segregation ratio of 69:100:49 corresponded with 1:2:1 Mendelian segregation ratio, indicating a goodness of fit for blast resistance gene loci. Figures 5 and 6 display some of the banding patterns obtained from foreground selection of BC_2_F_2_ plants. Table 3 shows the genetic composition of the 16 newly developed lines with their respective gene pyramids.

**Figure 5:**
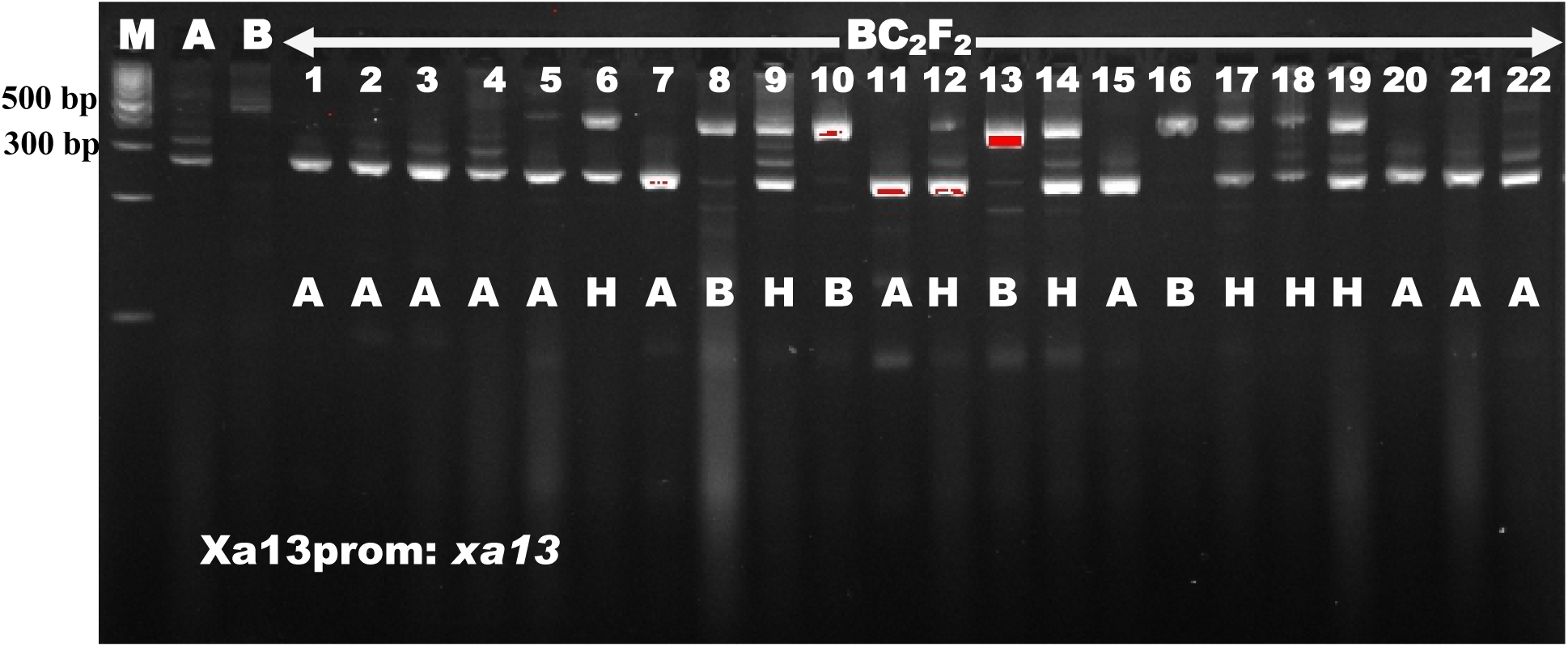
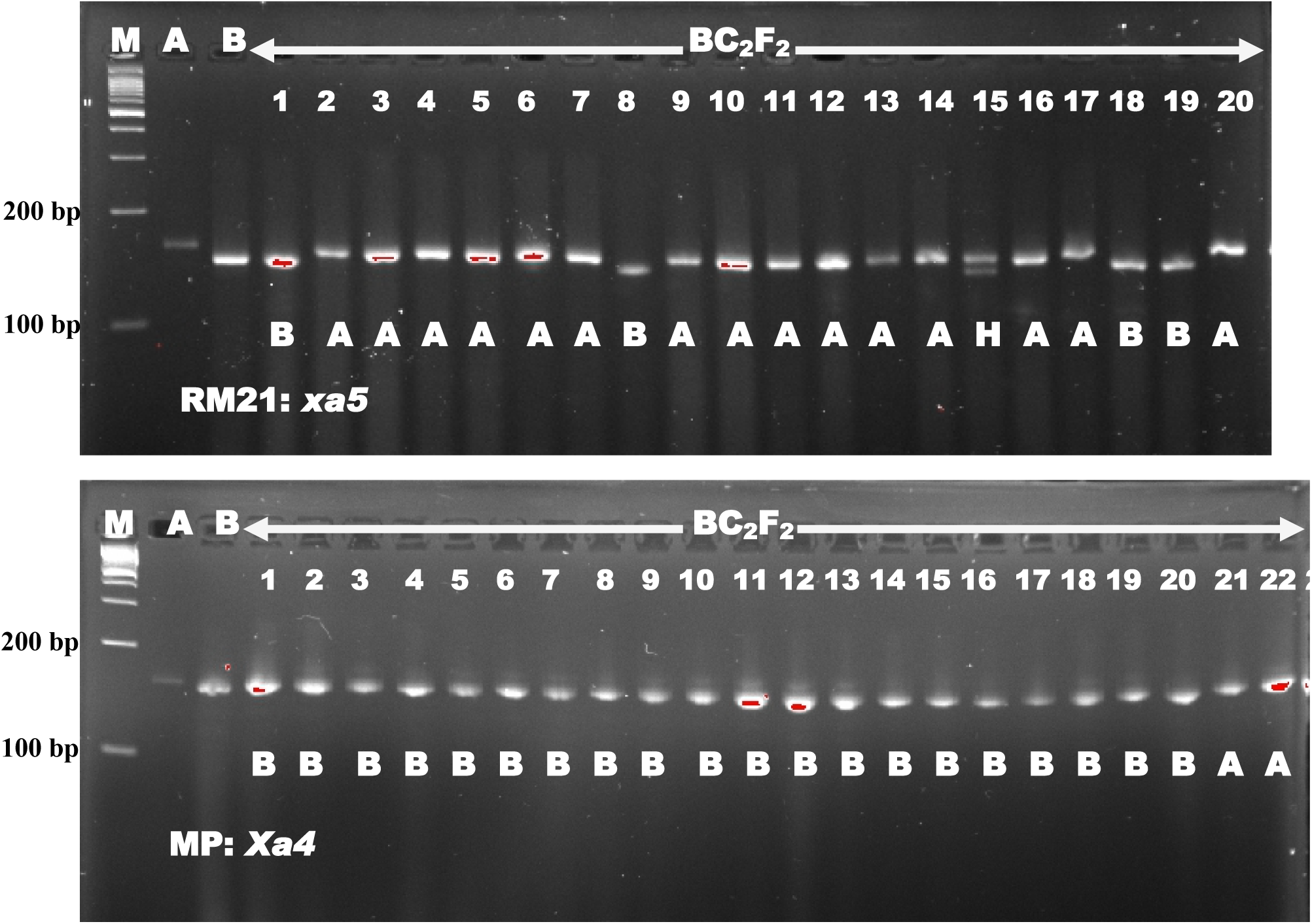
Genotyping and foreground marker-assisted selection of BC_2_F_2_ progenies using polymorphic tightly linked markers. Note: Only homozygous “B” allele for *Xoo* resistance genes were selected at BC_2_F_2._ M: 100bp DNA ladder

**Figure 6:**
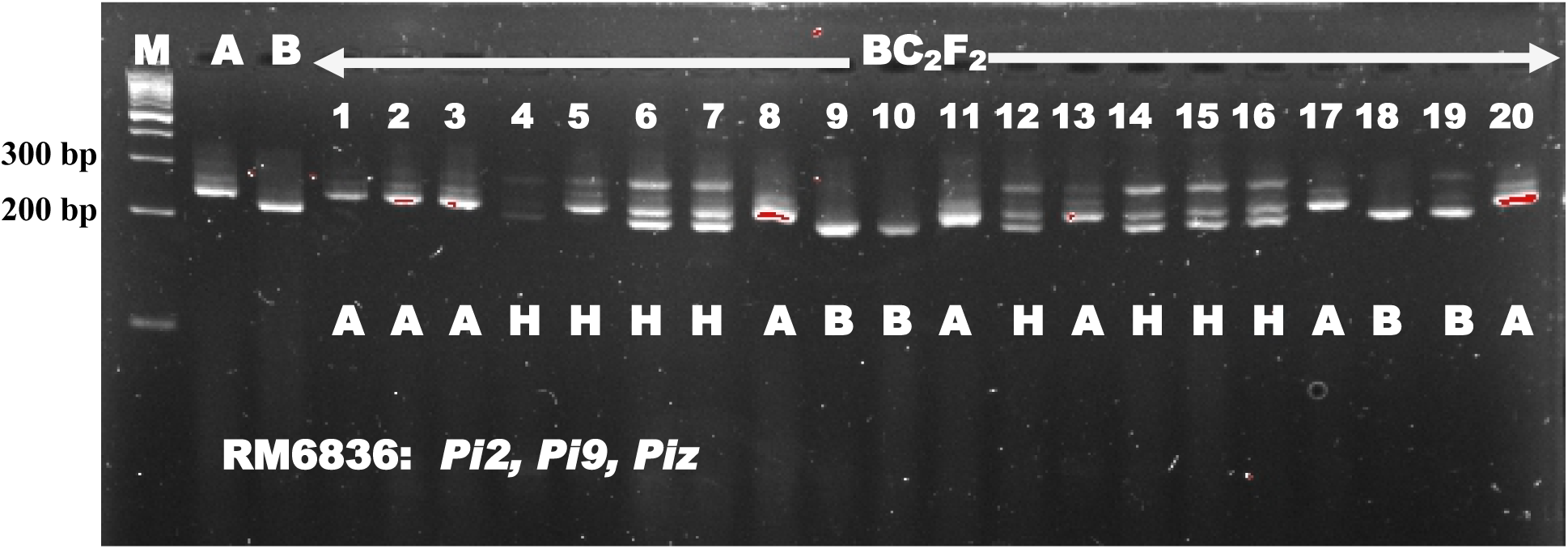
Genotyping and foreground marker-assisted selection of BC_2_F_2_ progenies using polymorphic tightly linked marker. Note: Only homozygous “A” allele for blast resistance genes were selected at BC_2_F_2._ M: 100bp DNA ladder

### Marker-assisted background selection of selected pyramided lines

In order to confirm that the pyramided lines maintained high yielding characteristic from the genetic background of Putra-1, the improved selected lines were molecularly screened using 79 polymorphic microsatellites markers. These 79 polymorphic microsatellites spread across the 12 chromosomes of rice covering the 1591 cM of the genome. The result in Table 2 shows the percentage RPGR and heterozygous segment of selected BC_2_F_2_ lines. The recurrent parent genome recovery obtained ranged from 93.2% in BC_2_F_2_-109 to 98.5% in BC_2_F_2_-4. The mean percentage recurrent parent genome recovery recorded in all the 16 selected lines across the 12 chromosomes was 95.9. The Chromosome-wise recurrent parent genome recovery (RPGR) of the 16 improved selected BC_2_F_2_ lines are shown in Figure 9. This result is an indication that the improved lines not only have been pyramided with *Xoo* and blast resistance genes, thereby incorporating a broad spectrum resistance against bacterial leaf blight and blast diseases but also maintained high yielding trait.

**Table 2:**
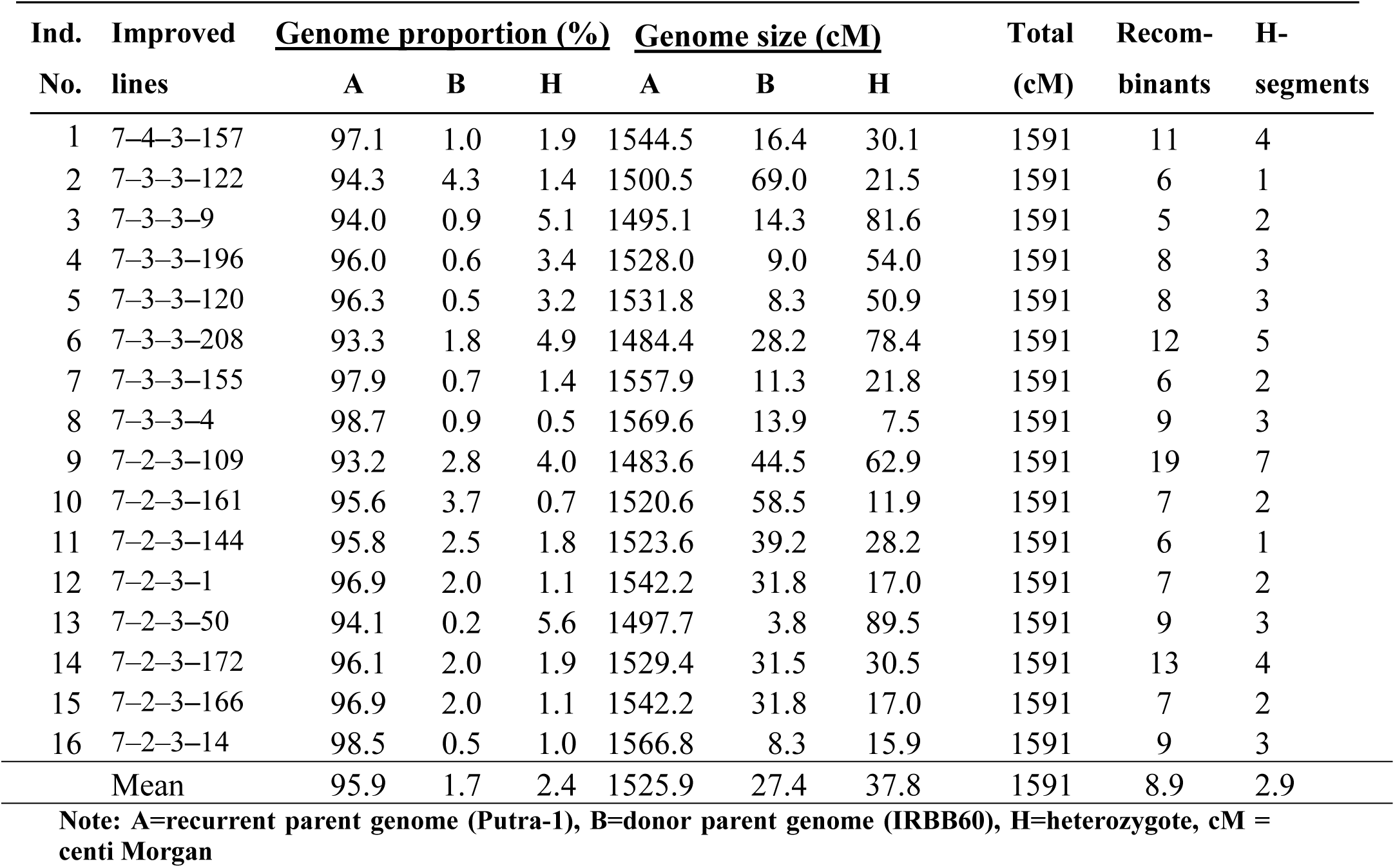
Percentage RPGR and heterozygous segment of selected BC_2_F_2_ lines

**Table 3:** Genetic composition of BC_2_F_2_ selected lines pyramided with multiple R-genes for broad spectrum and durable resistance

| s/n | Improved lines | Pyramided genes | Reaction to *Xoo* and *M. oryzae* pathogens |  |
| --- | --- | --- | --- | --- |
|  |  |  | Pathotype P7.7 | Pathotype P7.2 |
| 1 | 7-4-3-157 | *Xa21* + *xa13* + *xa5* + *Xa4* + *Pi2* + *Pi9* + *Piz* | HR | HR |
| 2 | 7-3-3-122 | *xa13* + *xa5* + *Xa4* + *Pi2* + *Pi9* + *Piz* | R | HR |
| 3 | 7-3-3-9 | *Xa21* + *xa13* + *Xa4* + *Pi2* + *Pi9* + *Piz* | R | HR |
| 4 | 7-3-3-196 | *Xa21* + *xa13* + *Xa4* + *Pi2* + *Pi9* + *Piz* | HR | HR |
| 5 | 7-3-3-120 | *xa13* + *xa5* + *Xa4* + *Pi2* + *Pi9* + *Piz* | R | HR |
| 6 | 7-3-3-208 | *Xa21* + *xa13* + *Xa4* + *Pi2* + *Pi9* + *Piz* | R | HR |
| 7 | 7-3-3-155 | *Xa21* + *xa13* + *Xa4* + *Pi2* + *Pi9* + *Piz* | R | HR |
| 8 | 7-3-3-4 | *Xa21* + *xa13* + *Xa4* + *Pi2* + *Pi9* + *Piz* | HR | HR |
| 9 | 7-2-3-109 | *xa5* + *Xa4* + *Pi2* + *Pi9* + *Piz* | R | HR |
| 10 | 7-2-3-161 | *xa5* + *Xa4* + *Pi2* + *Pi9* + *Piz* | R | R |
| 11 | 7-2-3-144 | *xa13* + *Xa4* + *Pi2* + *Pi9* + *Piz* | R | HR |
| 12 | 7-2-3-1 | *xa13* + *Xa4* + *Pi2* + *Pi9* + *Piz* | R | HR |
| 13 | 7-2-3-50 | *xa13* + *Xa4* + *Pi2* + *Pi9* + *Piz* | R | HR |
| 14 | 7-2-3-172 | *xa13* + *Xa4* + *Pi2* + *Pi9* + *Piz* | R | HR |
| 15 | 7-2-3-166 | *xa5* + *Xa4* + *Pi2* + *Pi9* + *Piz* | R | R |
| 16 | 7-2-3-14 | *xa13* + *Xa4* + *Pi2* + *Pi9* + *Piz* | R | HR |

### Phenotypic screening of parents and the hybrid F_1_ for bacterial leaf blight resistance

The result shows that percentage infection ranged from 42.86 to 63.70 (Table 4) in Putra-1 variety (the recurrent parent). The score of seven indicates that the Putra-1 variety was susceptible to BLB infection. The percentage infection in IRBB60 ranged from 0.00 to 3.08 while the mean percentage infection was recorded as 1.76. The mean percentage score of 1.76 and disease score of 1 showed that the IRBB60 was resistant to BLB infection caused by *Xoo*. In the F_1_ progeny, percentage infection recorded ranged from 4.24 to 10.91. The average percentage infection was 6.35 while the mean disease score was recorded as 1. These results indicated that the F_1_ progenies have been introgressed with *Xoo* resistance genes and were resistant to the BLB infection.

**Table 4:** Reaction of Putra-1 and IRBB60 parental varieties including the F_1_ progenies to *Xoo* isolate

| Genotype | % infection | Score | HR |
| --- | --- | --- | --- |
| Putra-1 | 52.61 | 7 | S |
| IRBB60 | 1.76 | 1 | R |
| F<sub>1</sub> hybrid | 6.35 | 1 | R |

#### 5.3.2 Phenotypic screening of selected improved BC_2_F_2_ lines and their parents for bacterial leaf blight resistance

No symptom (lesion) of bacterial leaf blight was observed on the leaves of IRBB60 plants. The donor parent for bacterial leaf blight resistance genes IRBB60 which carries the *Xoo* R-genes, displayed high level of resistance to bacterial leaf blight infection with a mean disease score of 0. The result shows that the donor parent IRBB60 has high level of resistance to *Xanthomonas oryzae* pv *oryzae* which is responsible for bacterial leaf blight infection. The recipient (recurrent) parent had a mean disease score of 7 indicating a susceptibility to bacterial leaf blight infection and this was evidenced by the symptoms observed on the plants. However, the disease score recorded for BC_2_F_2_ introgression lines ranged from 0 to 1 with a mean of 1. The results showed that the BC_2_F_2_ selected lines carried the *Xoo* R-genes and were resistant to the disease. The distribution of lesion length after inoculation with *Xanthomonas oryzae* pv *oryzae* in the parents and introgression lines are presented in Figure 7.

**Figure 7:**
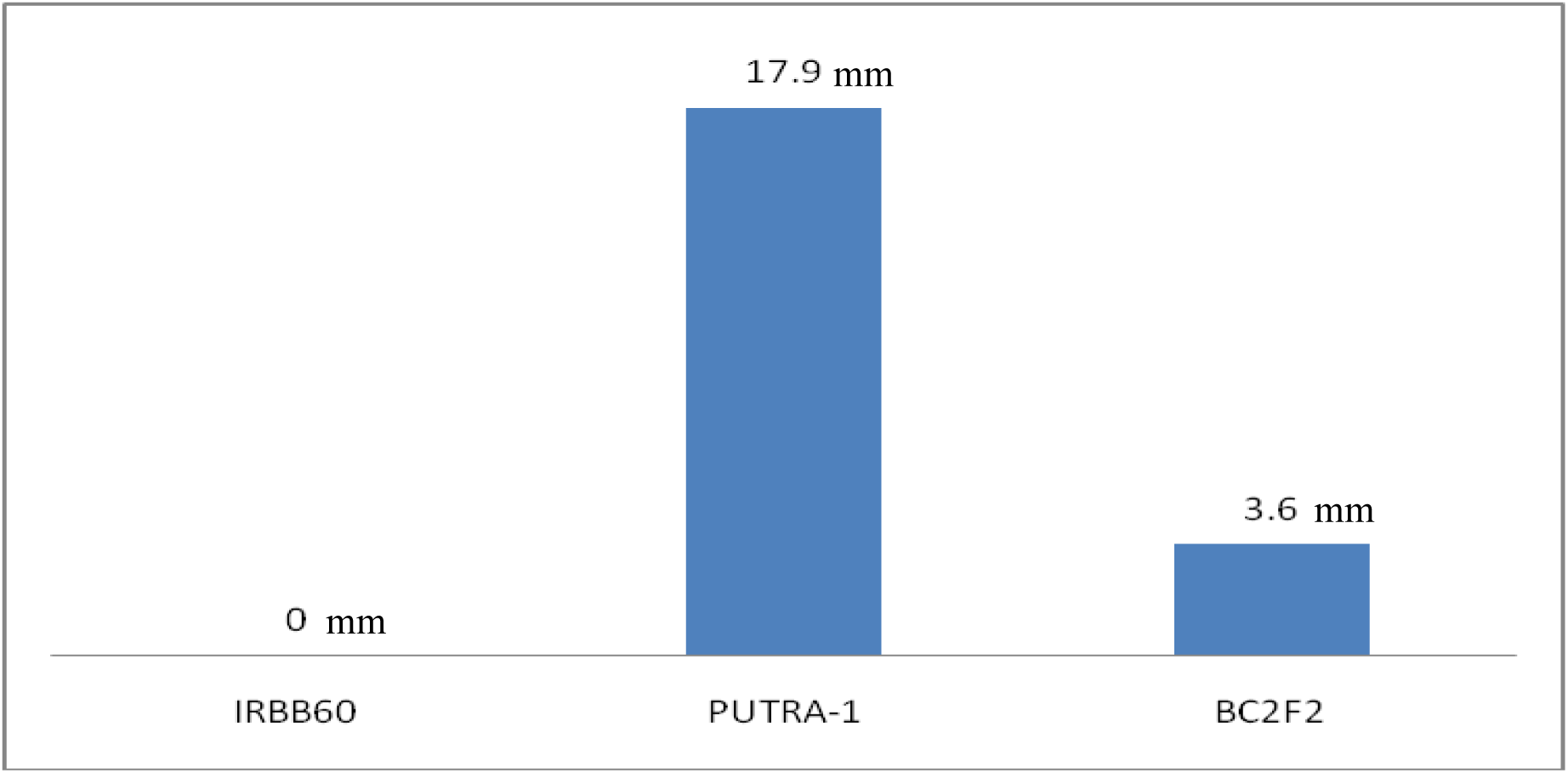
Distribution of lesion length after inoculation of *Xoo* pathotype P7.7 in parents and introgression lines of BC_2_F_2_ progenies.

#### 5.3.3 Phenotypic screening of BC_2_F_2_ lines and their parents for blast resistance

The donor parent for blast resistance genes Putra-1 which carries the *Pi* genes, displayed high level of resistance to rice blast with a disease score ranging from 0 to 1 while the mean score obtained was 0 indicating high resistance to blast infection. Incidentally, the other parent IRBB60 showed a moderate resistance to blast infection with a mean disease score of 3. The BC_2_F_2_ selected improved lines displayed resistance to blast disease with disease score ranging from 0 to 1 and a mean score of 1. The distribution of lesion length after challenging with *Magnaporthe oryza*e pathotype P7.2 in the parents and BC_2_F_2_ lines are presented in Figure 8.

**Figure 8:**
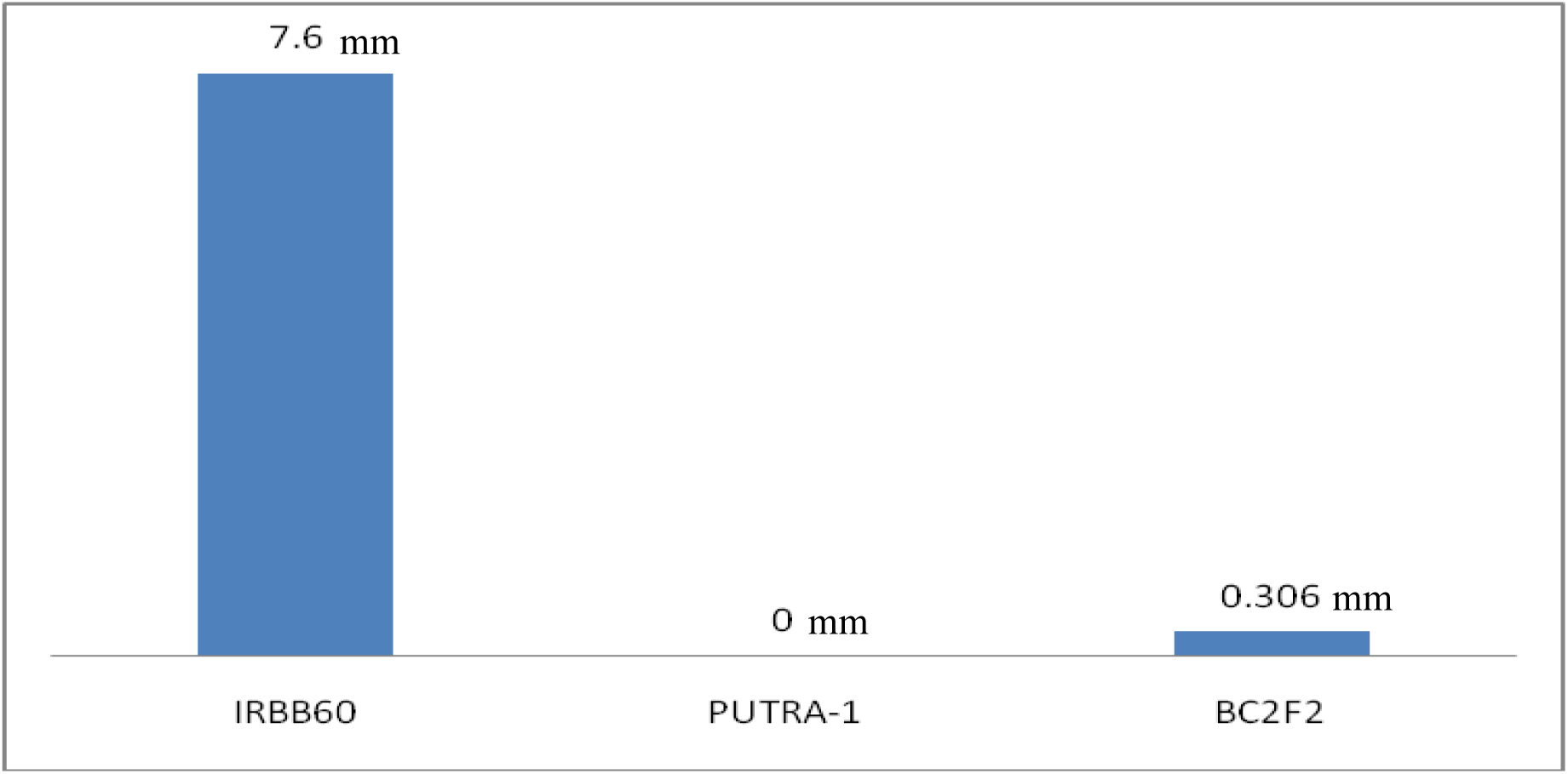
Distribution of lesion length after inoculation of *Magnaporthe oryzae* pathotype P7.2 in the parents and BC_2_F_2_ lines.

**Figure 9:**
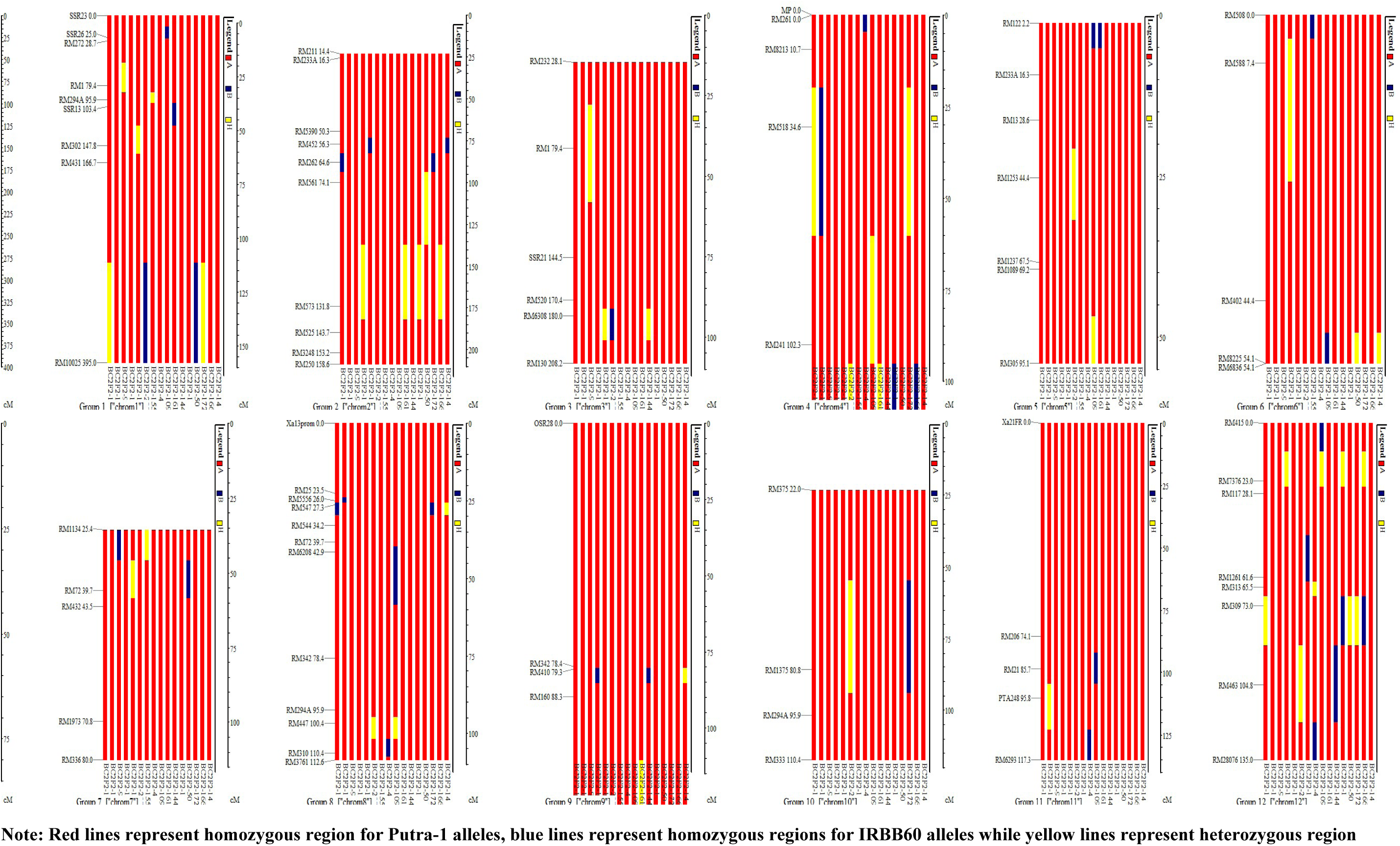
Chromosome-wise recurrent parent genome recovery (RPGR) of 16 improved lines.

The results of bacterial leaf blight and blast screening of BC_2_F_2_ progenies are as presented in Tables 5 and 6. The Chi-square result shows a significant difference (P<0.05) in segregation of resistance against *Xoo* in the single dominant gene model with regard to the expected and observed number of resistant and susceptible plants. The result implies inability of the plants to segregate according to the expected 3:1 ratio proposed by Mendel for single dominant genes. However, the Chi-square tests conducted on data obtained from segregation of resistance against blast in different gene models, viz; single gene model, two independent gene model and/or two locus interaction, revealed that the expected number of resistant and susceptible plants in the single dominant gene model’s segregation ratio did not differ significantly (P>0.05) from the observed number of resistant and susceptible plants, hence, they fitted to the 3:1 Mendelian segregation ratio. More so, the result in Table 6 shows there was a significant difference (P<0.05) in the observed vs expected ratio for the two independent gene model and epistasis effects both in bacterial leaf blight and blast. Therefore, the segregation ratio did not fit into the 9:3:3:1 and 15:1 ratio of two independent gene model and epistasis effects respectively. This is attributed to maker segregation distortion phenomenon.

**Table 5:** Observed and expected segregation ratios of resistant and susceptible plants in the BC_2_F_2_ generation challenged with pathotype P7.7 and P7.2 of *Xoo* and *Magnaporthe oryzae*

| **Reaction** | **Observed** | **Expected** | **Chi-square (3:1)** |
| --- | --- | --- | --- |
| **BLB** |  |  |  |
| Resistant | 206 | 165 | 10.19 |
| Susceptible | 14 | 55 | 30.56 |
| Total | 220 | 220 | 40.75 |
| **Blast** |  |  |  |
| Resistant | 155 | 165 | 0.63 |
| Susceptible | 65 | 55 | 1.89 |
| Total | 220 | 220 | 2.53 |
| **Df=1, $\chi^2(0.05,1)=3.84$** |  |  |  |

**Table 6:** Chi-square test for independent genes (9:3:3:1) and epistasis effects (15:1) for BLB and blast resistance in BC_2_F_2_ population inoculated with the most virulent pathotype P7.7 and P7.2 of *Xoo* and *Magnaporthe oryzae* respectively.

| Gene Model | Observed | | | | Expected | | | | $\chi^2$ values<br>(9:3:3:1) |
| --- | --- | --- | --- | --- | --- | --- | --- | --- | --- |
|  | R | MR | MS | S | R | MR | MS | S |  |
| BLB |  |  |  |  |  |  |  |  |  |
| Independent gene | 173 | 30 | 11 | 6 | 123.75 | 41.24 | 41.25 | 13.75 | 49.22 |
| Epistasis effect | 214 | — | — | 6 | 206.25 | — | — | 13.75 | 4.66 |
| Blast |  |  |  |  |  |  |  |  |  |
| Independent gene | 85 | 67 | 38 | 30 | 123.75 | 41.24 | 41.25 | 13.75 | 47.69 |
| Epistasis effect | 190 | — | — | 30 | 206.25 | — | — | 13.75 | 20.48 |

Where $\chi^2_{0.05, 3} = 7.81$ ; $\chi^2_{0.05, 1} = 3.84$
**Note:** R=Resistant; MR=Moderately resistant; MS=Moderately susceptible and S=Susceptible for two independent genes while R, MR and MS were considered as resistant and S as susceptible for epistasis effect.

### Agro-morphological performance of selected improved lines compared to the recurrent parent

The selected improved lines were evaluated based on twenty agro-morphological characteristics and the result was compared to the performance of their recurrent parent. The result showed that the newly improved lines differed significantly (P<0.05) from their recurrent parents in all the twenty agro-morphological traits evaluated except leaf area, leaf area index and number of unfilled grains which were statistically the same (Table 7). Plant height was significantly reduced to 110.65 cm compared to the taller height of 116.50 cm recorded for their recurrent parent. Also, days to flowering and maturity of the selected improved lines were significantly reduced by 10 and 15 days, respectively, compared to what was obtained in the recurrent parents. However, there was significant increase in other major yield component traits such as number of panicles per hill, number of tillers, panicle length, number of leaves, total number of grains per panicle, 1000-grain weight and total grain weight per hill. The yield per hectare was also significantly (P<0.05) increased by 0.37 tonnes per hectare as compared to the yield of the recurrent parent (Table 7).

**Table 7:** Descriptive *t*-test and co-efficient of variation for mean comparison between the selected improved BC_2_F_2_ lines and their recurrent (recipient) parent using independent *t*-test

| LINES | PH<br>(cm) | FLL<br>(cm) | FLW<br>(cm) | LA<br>(cm<sup>2</sup>) | LAI | FLWR | NP/H<br>(no.) | DF<br>(days) | DM<br>(days) | NL<br>(no.) | NT<br>(no.) | PL<br>(cm) | TNG/P<br>(no.) | NUFG<br>(no.) | 1000GW<br>(g) | TGW/H<br>(g) | SL<br>(cm) | SW<br>(cm) | SLWR | Y/HA<br>(t/ha) |
| --- | --- | --- | --- | --- | --- | --- | --- | --- | --- | --- | --- | --- | --- | --- | --- | --- | --- | --- | --- | --- |
| BC<sub>2</sub>F<sub>2</sub> | 110.65<sup>a</sup> | 36.85<sup>a</sup> | 2.81<sup>a</sup> | 103.62<sup>a</sup> | 0.17<sup>a</sup> | 13.14<sup>a</sup> | 15.94<sup>a</sup> | 75.75<sup>a</sup> | 105.75<sup>a</sup> | 80.31<sup>a</sup> | 16.06<sup>a</sup> | 32.64<sup>a</sup> | 177.56<sup>a</sup> | 42.81<sup>a</sup> | 79.45<sup>a</sup> | 52.74<sup>a</sup> | 0.95<sup>a</sup> | 0.25<sup>a</sup> | 3.86<sup>a</sup> | 8.44<sup>a</sup> |
| SE | ±1.13 | ±1.35 | ±0.04 | ±4.15 | ±0.01 | ±0.48 | ±0.84 | ±0.66 | ±0.66 | ±4.44 | ±0.89 | ±0.62 | ±7.40 | ±5.30 | ±0.54 | ±2.32 | ±0.02 | ±0.00 | ±0.08 | ±0.37 |
| Recurrent<br>parent | 116.50<sup>b</sup> | 33.80<sup>b</sup> | 2.70<sup>b</sup> | 68.28<sup>a</sup> | 0.11<sup>a</sup> | 12.56<sup>b</sup> | 15.00<sup>b</sup> | 85.67<sup>b</sup> | 120.67<sup>b</sup> | 75.00<sup>b</sup> | 15.00<sup>b</sup> | 31.83<sup>b</sup> | 148.00<sup>b</sup> | 115.33<sup>a</sup> | 75.53<sup>b</sup> | 50.41<sup>b</sup> | 0.97<sup>b</sup> | 0.25<sup>b</sup> | 3.92<sup>b</sup> | 8.07<sup>b</sup> |
| SE | ±1.40 | ±1.89 | ±0.06 | ±2.36 | ±0.00 | ±0.97 | ±0.58 | ±1.45 | ±0.88 | ±2.89 | ±0.58 | ±1.05 | ±3.61 | ±7.22 | ±0.75 | ±1.86 | ±0.02 | ±0.00 | ±0.11 | ±0.30 |
| Independent<br>t-test | 38.83 | 23.16 | 51.82 | 4.86 | 4.94 | 44.61 | 33.00 | 16.27 | 15.18 | 29.24 | 29.24 | 28.19 | 11.01 | 2.18 | 23.85 | 44.35 | 78.59 | 110.11 | 135.89 | 44.89 |
| P<0.05 | 0.0164 | 0.0275 | 0.0123 | 0.1291 | 0.1271 | 0.0143 | 0.0193 | 0.0391 | 0.0419 | 0.0218 | 0.0218 | 0.0226 | 0.0576 | 0.2737 | 0.0267 | 0.0144 | 0.0081 | 0.0058 | 0.0047 | 0.0142 |
| CV | 3.64 | 6.11 | 2.73 | 29.07 | 28.61 | 3.17 | 4.29 | 8.69 | 9.32 | 4.84 | 4.84 | 5.03 | 12.84 | 64.85 | 5.93 | 3.19 | 1.80 | 1.28 | 1.04 | 3.15 |

**Note:** PH=plant height, FLL=flag leaf length, FLW=flag leaf width, LA=leaf area, LAI=leaf area index, FLWR=flag-leaf length:width ratio, NP/H=number of panicles per hill, DF=days to 50% flowering, DM=days to maturity, NL=number of leaves, NT=number of tillers, PL=panicle length, TNG/P=total number of grains per panicle, NUGF=number of unfilled grain per panicle, 1000GW=one thousand grain weight, TGW/H=total grain weight per hill, SL=seed length, SW=seed width, SLWR=seed length:width ratio, Y/HA=yield per hectare, CV=co-efficient of variation.

## Discussion

In this study, eight polymorphic tightly linked foreground markers were used to confirm the pyramiding of seven *Xoo* and blast resistance genes in the newly improved lines. The four *Xoo* resistance genes (*Xa4 + xa5* + *xa13* + *Xa21*) were introgressed from IRBB60 variety into the Malaysia elite variety Putra-1 with genetic background of three *Pi* genes, namely; *Pi9, Pi2* and *Piz* leading to development of lines with seven gene pyramid (*Xa4+xa5*+*xa13*+*Xa21+Pi9+Pi2*+*Piz*) for broad spectrum resistance. Pyramiding may involve incorporating genes from two or more parents. For example, Pradhan *et al*. [4] pyramided three bacterial leaf blight (*xa5* + *xa13* + *Xa21*) resistance genes for broad-spectrum resistance in deep water rice variety, Jalmagna. Three major genes (*Pi1, Piz-5* and *Pita*) were pyramided by Hittalmani *et al.* [24] using RFLP markers. Hittalmani et al. [24] and Castro et al. [25] combined genes originating from three parents for rice blast and stripe rust in barley, respectively. Gene pyramiding is difficult to achieve using conventional breeding alone because of linkage with some undesirable traits that is very difficult to break even after repeated backcrossings [26]. When two or more genes are introgressed, phenotypic evaluation is unable to distinguish the effect of individual gene precisely since each gene confers resistance to and combats multiple races of the pathogen. Moreover, in the presence of a dominant and a recessive allele, the effect of the recessive gene is masked. The advent and easy availability of molecular markers closely associated with each of the resistance genes makes identification of plants with multiple genes possible. Earlier efforts to generate three-gene pyramids in the backgrounds of rice cultivars PR106 [27], Pusa Basmati [28] and Samba Mahsuri by the collaborative work of the Directorate of Rice Research, Hyderabad, India (DRR) and the Centre for Cellular and Molecular Biology, Hyderabad, India (CCMB) have been successful [29].

Studies conducted to identify the best gene combinations conferring broad-spectrum resistance showed that the four-gene combination was the most effective and did not show any sign of breakdown of resistance to various strains of the BLB pathogen from different parts of the country [30]. Shanty et al. [31] worked on the pyramiding of four BLB resistance genes *Xa4, xa5*, *xa13* and *Xa21* through marker-assisted selection (MAS) in the BLB susceptible high yielding rice cv. Mahsuri and parental lines of hybrid rice, maintainers IR58025B and Pusa 6B and the restorer lines KMR3 and PRR78 simultaneously. The results showed increased and broad spectrum resistance to the pathogen populations from different parts of India. The work was the first successful example of the use of molecular markers in foreground selection in conjunction with conventional breeding for the simultaneous introgression of genes of interest into multiple backgrounds [31]. Popular plant varieties, adopted in large-scale production, may have some undesirable traits. These cultivars are generally high-yielding but may be highly susceptible to the attack of a certain disease or insect. In some cases, these are of poor grain quality, and thus command lower prices in the market. Also, long-term cultivation of single variety with wide adoption can lead to a shift in race or biotype frequency and ultimately cause resistance breakdown [32].

One way to delay loss of resistance is called gene pyramiding or gene stacking [33, 27]. Thus, to further extend the use of these popular varieties, new traits or genes are being introduced through a series of backcrossing [34]. However, pyramiding is difficult or impossible using the conventional approach due to masking and/or epistatic effects of the genes being combined [35]. Moreover, some genes have similar reactions to two or more races or biotypes; thus they prove rather difficult to identify and transfer through conventional approaches [27]. Most of the rice varieties in the Philippines have *Xa4* and *xa5* genes for bacterial leaf blight resistance and some have no resistance at all [36]. With the identification of new genes like *Xa21, Xa22* and *Xa23* or the *Om* gene from *Oryza minuta,* varieties developed earlier can be improved through the incorporation of these new important genes [37]. Several genes for BLB have been mapped and tagged using DNA markers and these can be explored in improving resistance [38]. Among the more than two dozen genes reported [39, 40, 41, 42, 43], eight genes (*Xa3, Xa4, xa5, Xa7, xa8, Xa10, xa13* and *Xa21*) have been transferred or pyramided in common genetic background using marker-aided selection [44, 33, 27, 45, 35, 46]. Lines with single or combinations of genes have been identified using different DNA markers closely linked to the genes. Classical transfer of new genes is being done through a series of backcrossing and selection, and this procedure needs a longer time and requires large resources. The use of DNA as markers for selection has streamlined and facilitated the whole process even without inoculation [37].

In this study, marker segregation distortion was observed in *Xoo* resistance genes at BC_2_F_2_. The markers at those loci failed to show a goodness of fit to the expected Mendelian ratios. A study conducted by Jin et al. [47] showed that in 300 F_2_ populations evaluated, the foreground marker segregation for the *BADH2* gene did not correspond with the expected 1:2:1 Mendelian segregation ratio. Also, Amarawathi et al. [48] reported non-fitness to the 1:2:1 ratio of the nksbad2 marker, hence segregation distortion in their RIL individuals. Lau [49] also observed a segregation ratio of 69:46:44 which failed to fit into the 1:2:1 ratio and the result was attributed to segregation distortion of foreground markers. In the same vein, Muthusamy et al. [50] reported distortion of foreground marker segregation which was observed in all the backcross stages performed in maize and this was without regard to the size of population evaluated. Therefore, population size of BC_2_F_2_ progenies could not be taken as a factor responsible for the foreground markers segregation distortion observed in this study. The BC_1_F_1_ and BC_2_F_1_ both fitted into the Mendel’s single gene segregation ratio despite having varying population sizes. The distortion of foreground markers segregation observed at *Xoo* resistance locus could be attributed to variation in environmental factors. Bing et al. [51] and Xu et al. [52] all reported that distortion of marker segregation could be as a result of environmental issues. Similarly, studies conducted by Muthusamy et al. [50] showed that the distortion in foreground marker segregation in maize backcrossings was due to backcrossings performed during winter season. Wang et al. [53] discovered distortion of marker segregation particularly in F_2_ population which were grown at a particular altitude, but when grown at other altitudes, marker segregation distortion was not detected. There is, therefore, every likelihood that the markers segregation distortion observed in this study was due to environmental effects as the BC_2_F_2_ progenies were planted at varied weather conditions compared to BC_1_F_1_ and BC_2_F_1_ in 2019 and 2018 as obtainable in Malaysia’s agro-ecology. Notwithstanding, the functional markers used in this study facilitated the identification of the genotypes with the targeted *Xoo* resistance genes in addition to the already established background genetic make-up of blast resistance in Putra-1.

This Chi-square result obtained for blast resistance loci was in line with the result reported by Tanweer et al. [54] who recorded foreground marker segregation ratio of 53:106:41 and 55:107:38 for RM208 and RM206 respectively which fitted into the 1:2:1 segregation ratio expected from Mendel studies. Also, Miah et al. [15] reported a segregation ratio of 52:127:74 (RM8225) and 54:132:67 (RM6836) which did not differ significantly from the expected Mendelian ratio of 1:2:1. It is important to confirm the introgression of target gene by phenotyping in the developed cross mainly at individual level as individual phenotypic performance is a good indicator of the genotype provided that genes have major effect on phenotypic expression and the phenotyping error is less [55]. Phenotypic selection for desirable agro-morphological traits has always been done along with backcross selection [23]. On the other hand, Young and Tanksley [56] proposed that the genotypic selection which was later referred to as background selection [57] involved screening the allelic pedigree throughout the genome with the aid of markers. Singh et al. [58] reported that foreground selection for target genes followed by stringent phenotyping gives superior yield.

The high recurrent parent genome recovery in this study could be as a result of the stringent phenotypic selection done at every backcross generation and selfing. This strategy reduced both time and economic resources input. Singh et al. [59] also reported that phenotypic and marker-assisted selections increase the RPGR. The result obtained in this present study was in contrast with the findings made by Sabu et al. [60] and Martinez et al. [61] who found no significant difference in grain yield between the parental cultivar and its progenies. Hossain et al. [62] noted that number of panicles is the result of productive tillers which survived to produce panicles. The significant increase in yield contributing traits such as number tillers, number of panicles, number of leaves, etc., recorded in this present study could have resulted to the significant increase in grain yield per hectare obtained. Dutta et al. [63] and Kusutani et al. [64] both reported that higher grain yield in rice genotypes is associated with number of productive tillers per hill and number of grains per panicle. Similarly, seed length and width are essential quantifiable characteristics that are closely related to the external physical quality of rice [65, 66]. Juliano [67] reported that seed length and width determines the grain/seed shape of rice. There was a significant difference in the seed length and width of the newly improved selected rice lines compared to their original recurrent parent. However, both lines have slender seed shape (seed length:width ratio >3.0). Shi and Zhu [68](1996) reported that seed shape is concurrently controlled by cytoplasmic and maternal genes as well as triploid endosperm

## Conclusion

The four *Xoo* R-gene from IRBB60 were successfully introgressed into the backcross progenies and fixed in plants that had homozygous *Xa-*genes similar to IRBB60 and homozygous *Pi*-genes similar to Putra-1 in the BC_2_F_2_ generation. Based on the MABB method and combined phenotypic selection strategy adopted in this breeding programme, 16 progenies from the BC_2_F_2_ generation were selected as advanced backcross rice lines with characteristic high yielding, resistance to bacterial leaf blight and blast diseases. The result obtained from this study reveals that resistance to *Xoo* pathotype P7.7 in IRBB60 was neither due to two independent gene action nor epistasis but most likely due to single nuclear gene action. The result also clearly showed that the inheritance of blast resistance to pathotype P7.2 was mainly due to single gene action. More so, the pyramiding of seven resistance genes in the newly developed lines shows the presence of horizontal resistance. It is suggested that only the existence of horizontal resistance along with the vertical component could help varieties with enhanced and sustainable resistance to bacterial leaf blight and blast. Therefore, the pyramiding of seven BLB and blast resistance genes (*Xa4+xa5*+*xa13*+*Xa21+Pi9+Pi2*+*Piz*) in the newly developed lines confers broad spectrum resistance to bacterial leaf blight and blast diseases. The newly developed lines are recommended as new varieties suitable for commercial cultivation in Malaysia.

## Acknowledgements

The authors would like to thank all researchers and colleagues whose published works were useful as reference during the study. This study was supported by the Higher Institution Centre of Excellence (HiCoE) Research Grant (Vot number 6369105), Ministry of Education, Malaysia. We express our appreciation for the fund received from HiCoE for research activities of improvement of rice crop against diseases.

